# Evaluation of Seed or Grain Storage Technologies from Indian Farming Societies Using A Hybrid and Multi-Attribute Approach

**DOI:** 10.1101/2020.06.25.172189

**Authors:** Aniruddha Maity, Anirban Mukherjee, Mrinmoy Ray, Pragati Pramanik, Sanjay S Parmar, N Manjunatha, Sanjay Kumar, Vinod Kumar Wasnik, Kumari Shubha

## Abstract

Soon after the introduction of green revolution, a parallel increment in percent food grain production and losses has been reported in India, which impedes the mission of achieving food security in India. In parts, among the major causes of these losses are the reluctance among farmers, especially of small holding, towards adoption of scientific storage methods/technologies and inability of the national agencies to meet the challenges imposed by the supply of surplus grains to them by the farmers. Although sets of traditional and improved grain storage technologies exist among Indian farming communities, a controversial account about their effectiveness and performance is available in the literature which negatively impacts the endeavour of bringing rapid developmental in the farming society. In a developing nation like India, a large section of farmers is often either unaware of the modern technologies or cannot logistically access them. They often rely on the traditional or semi-modern technologies without having a comprehensive account of the benefit and economic feasibility of the modern technologies. This impedes the vision of rapid development of the farmers and the process of infusing emerging technologies in the society. In order to develop an informative argument about the performance of the existing technologies, a nearly comprehensive study has been undertaken to compare the different traditional and improved grain storage structures used by Indian farmers. Both qualitative and quantitative parameters were compared individually and simultaneously to achieve their effect on performance individually and in unison. Another objective of the study was to test the applicability of hybrid and multi-attribute approach, based on Analytic Hierarchy Process (AHP) and the Technique for Order Preference by Similarity to Ideal Solution (TOPSIS) for the performance evaluation and ranking of the attributes and structures. Finally, based on analysis and ideas borrowed out of literature a set of theoretical and conceptual guidelines to assist improvement in these structures was brought in. The study will help understand the status of grain storage technologies in the developing, resource-poor society and also increase the adoption and access of farmers to better post-harvest seed storage technologies.

## Introduction

India has been among the primary beneficiaries of the Green Revolution (GR) (Pingali 2012), and since then has witnessed an unprecedented improvement in productivity of major cereal grains. For developing nations, like India, these developments have been instrumental in poverty reduction and food security, mostly by managing and distributing the surplus grains from resourceful areas to starved, marginal section of the society (Malek et al. 2017, Singh 2010). However, in India, during past few decades, a nationwide inadequacy in preventing and reducing post-harvest losses of stored grains or seeds have been reported (Mishra et al. 2012, Nagpal and Kumar 2012). While grain is the major source of food, seed is the basic input of agriculture to keep the food production system alive. Grain as well as seed productions follow a long chain of agronomic activities followed by biological processes in the plants and thereafter human intervention for its processing, storage and distribution (Maity and Pramanik 2013, Maity and Chakrabarty 2013). If any of the step is exposed to adverse weather conditions they lose their value and sometimes they become unusable (Maity et al. 2016)

According to the government reports available from 2008 to 2011, about 17,423 tonne of wheat worth $ 4.7 million had gone wasted (CAG 2013). Many similar cases have also been cited in news reports, for example, a nationally recognized news media reported a loss of about 1,068 million tons of food grains in Food Corporation of India (FCI) depots (Hindustan Times 2010). Similarly, according to the news agency Hindu, FCI admitted a loss of 1.945 million tons of food grains from 2005 to 2013 in their depots (The Hindu 2014). As per May 2009 price index, the total post-harvest losses within the country was about $ 6,830 million (CIPHET 2010) of which about $16,000 was for food grains (ASSOCHAM 2013). In these reports, evaluation of the causes of these storage losses were summarised in the form of poor management, lack of adequate facilities and large budgetary deficits. However, as opined by of different experts, like in other developing countries the government agencies of India and their policies have failed to fill the gap of the uneven social and spatial distribution of GR benefits, among favourable and unfavourable marginal and less favoured areas (herein after we will use the term “marginal”) which has led to these situations (Malek et al. 2017). Accordingly, the grain management policies adopted on the basis of subsidies have risked the financial health of these agencies, which is more noticeable in recent past due to fuel price hike and lack of funding from private partners (Malek et al. 2017, Taylan et al. 2015). In addition, forced selling of the surplus produce during harvest and reluctance among marginal farmers to increase and improve their own storage capacities have overburdened the national agencies, which are mandated to work even beyond their capacities (Singh 2010, Ganesh Kumar 1992). Therefore, concerns have been raised as the expected rise in agriculture productivity owing to introduction and use of more hybrid crops, would further complicate the situation, if new and innovative measure to reduce storage losses are not adopted at this stage (Gao and Hailu 2013, Singh 2010).

In India, in the context of technology, following traditional ‘pipeline approach’ (Malek et al. 2017, during last few decades several organizations and institutions including-governmental; national/international; private sector; non-governmental organizations (NGOs) etc. have undertaken a lot of work and studies related to development of new technology for farmers, for references see (Kumar and Kalita 2017, Gao and Hailu 2013, Parfitt et al. 2010). Some of these solutions were designed to cover specific commodities (Naveenaa et al. 2017), whereas others were found either impractical to or unadopted by the small farmers (Gao and Hailu 2013). In response to these insights, several experts and international organizations have favoured the development and adoption of demand-driven ‘holistic’ approach instead of traditional ‘pipeline approach’ (Malek et al. 2017, Singh 2010). Though details for pursuing and implementation of any of these “holistic” approaches vary with experts, but all have stressed the active participation of farmers and have advocated the capacity building for marginal farmers in their particular ecological and institutional environments (Malek et al. 2017, Rajalahti 2012).

In India, marginal farmers represent about 80% of the agriculture community, share around 83% of the total land holdings and contribute around 70% of the annual crop production of which 52% is through cereals production (Dev 2012). Therefore, the future food security of the country substantially depends on the performance of subsistence farming and reduction of seed or grain losses at their levels during storage must be perceived among the major priorities.

In India, two broad categories of storage structures are predominantly used by the farmers depending on their resources: traditional or indigenous structures and new or modern storage structures (Karthikeyan et al. 2009, Mann et al. 2016). The traditional storage facilities include various structures designed and mastered by ancient and/or local farming communities across the country. Their design and application vary with agroclimatic conditions and are made from a variety of locally available materials (Naveenaa et al. 2017). The modern storage structures include storage facilities developed and designed by different research and development organizations and commercial suppliers across the country (Mann et al. 2016).

Several previous studies have compared both traditional and modern structures, in terms of their efficiency to provide safety to stored commodity, and a sheer conflict could be observed among them. Arguments, in support of modern structures claims their better functioning in terms of temperature, humidity control and their efficient control of rodents and pests (Kumar and Kalita 217, Naveenaa et al. 2017). Similarly, for traditional structures, it has been claimed, that since these structures have evolved around certain specific agroclimatic conditions, they are suitable for storing grains in their original conditions only (Mann et al. 2016, Karthikeyan et al. 2009, Lamichaney et al. 2019). In addition, they have also reported their benefits, in terms of efficiency, associated cost and eco-friendly nature. Most of the post-green revolution cultivars grown today are often more susceptible to storage losses and for the sake of arguments, it is also true that traditional methods were developed around more robust local landraces (Ganesh Kumar 1992). Despite all efforts, in the background of all these arguments, it is also true, that due to various reasons adoption of modern structures has mostly been limited and mostly the subsistence farmers lack enough seeds to support their different needs (Naveenaa et al. 2017).

Usually, the comparison of different seed storage practices is assessed using traditional techniques that involve quantitative attributes like construction material and variations observed in physio-chemical and biological properties of the stored commodity during a specified period of time. However, in practice, selection of an appropriate storage structure by farmers depends on several qualitative attributes, like their socio-economic status, cultural ethnicity, future crop expectations, availability of transportation etc. (Mann et al. 2016, Karthikeyan et al. 2009. Hence, any evaluation process must simultaneously integrate all these quantitative and qualitative attributes during assessment.

Decision based on the traditional operations research models mostly include the risks of ignoring the effect of different qualitative factors, on the other hand attempting to include these qualitative factors may risk the decision being unscientific or ad-hoc nature of the methods. Additionally, the unavailability of precise data also limits the utility of traditional operations research models in analysing agricultural decisions (Alphonce 1997). Unfortunately, current agricultural statistics of India sometimes lacks quality, precision and reliability (Dev 2012, Parfitt et al. 2010).

With this background, we aimed to take a closer look at the post-harvest storage technologies available and used by Indian farmers, in a view to identify the gaps and possibility of improvement therein. Also, we used the Technique for Order Preference by Similarity to Ideal Solution (TOPSIS) and the Analytic Hierarchy Process (AHP) based hybrid multi-attribute decision model in present study to overcome existing complications with other tools.

## Material and Methods

For any multi-criteria evaluation process, the primary step is always to determine the attributes or parameters that would be included in the study followed by evaluation of these parameters using a set of defined techniques. In the present study, following activities were carried out to achieve the objectives.

### Characterization of Storage Structures

The range of structures available and used by Indian farmers varies geographically. It is diverse in design and continuously modified across generations based on use and experience by our ancestors or by the scientist based on scientific experiments. To get a comprehensive coverage, structures were selected on the basis of their design, crop coverages and perceived benefits, cost involved etc. through literature and experts’ judgments. In this study, eight traditional and six modern seed storage structures were considered based on their popularity and wide usability in India. The traditional seed storage structures were *kothi, gummi, khalda, sandook, kuthar, matka*, brick structure and underground structure (Fig 1a and b) whereas modern storage structures were Pusa bin, PAU bin, Hapur bin, Baked clay bin, Metal bins, Udaipur bin (Fig 1c and d).

**Figure 1.**
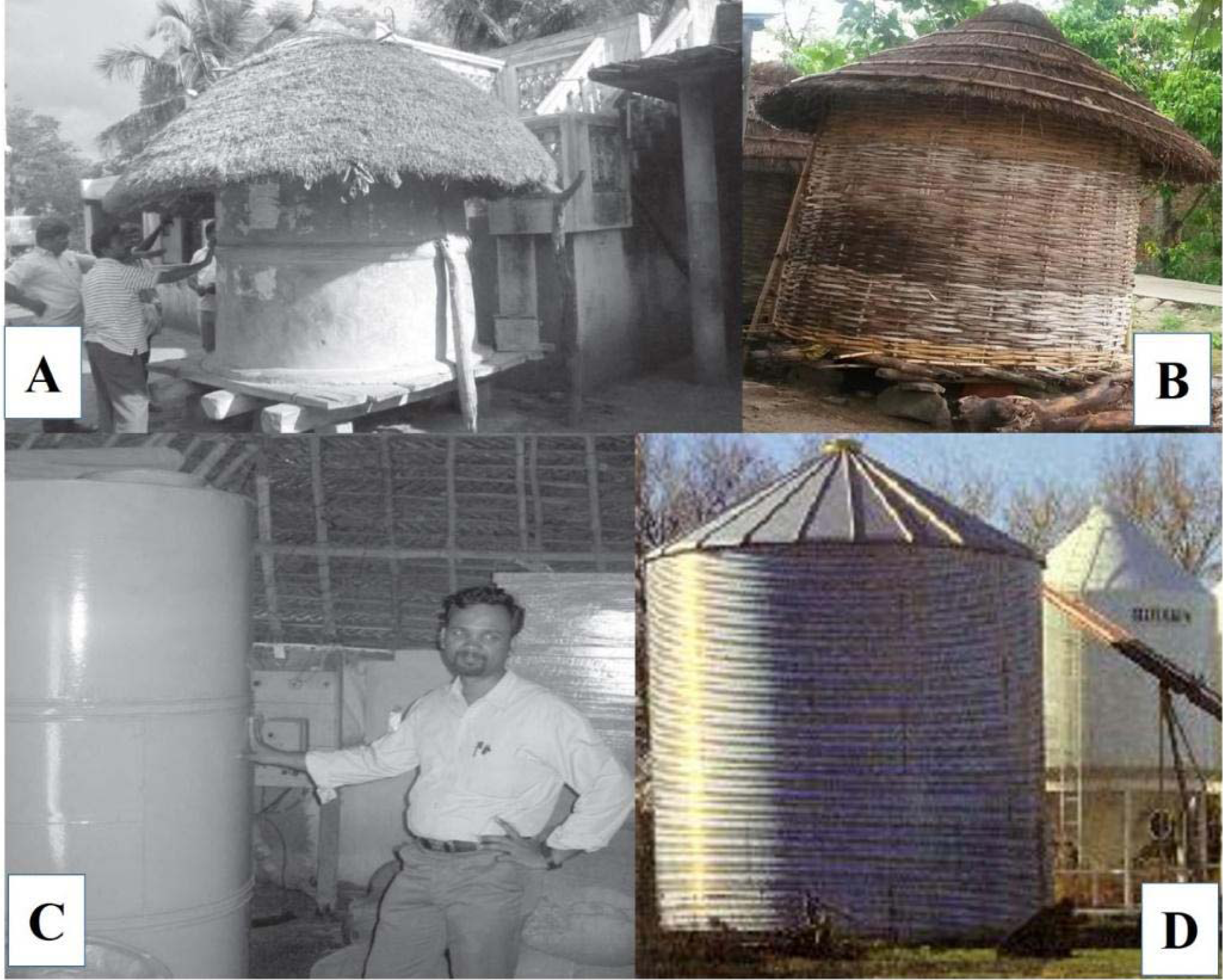
Traditional storage structure (Photo courtesy: (a) www.ciks.org; (b) Dr. P. K. Pathak, IGFRI) and modern storage structure (Photo courtesy: (c) www.ciks.org; (d) Canada Plan Service)

The features of traditional as well as modern seed storage structures were identified based on the selected parameters through the literature review, expert judgments and personal observations (Table 1). For performance evaluation and comparison, different parameters essential for effective storage of seeds/grains were selected. Overall ten important parameters viz. temperature control, humidity control, efficiency of plant protestants used, protection from rodents, strength of material used, capacity to store, life span of structure, life span of seed, handling and cost were identified through literature review and experts’ suggestion and were prioritized through analytical hierarchy process (AHP). These structures are used by different community located in different geographical areas for different crops. Majority of the community are using maximum two or three seed storage structures and can share their experience. Getting the primary and first-hand data by the same users for all the seed storage structures and then comparison was found difficult. Therefore, a group of sixty experts having more than ten years of experience in seed related research or field survey across the country were selected. A questioner was prepared and sent to all of them with standard instructions through e-mail. Based on these, the criteria for measurement were developed. Each and every storage structure was scored based on the criteria (Table 2).

**Table 1a.** Details of the traditional seed or grain storage structures used in the study.

|  | **Parameters** | **Kothi** | **Gummi** | **Khalda** | **Sandook** | **Kuthar** | **Matka** | **Brick structure** | **Underground structure** |
| --- | --- | --- | --- | --- | --- | --- | --- | --- | --- |
| 1 | Strength of material | Wood, bamboo, soil | Bamboo | Leather | Wood | Wood (Hut) | Earthen pot | Brick + cement | Mud, straw and stone |
| 2 | Capacity | 3-8q | 3-5q | 15-25kg | 7-8q | 22-25q | 15-20kg | 22-25q | 100-200q |
| 3 | Life span of structure | 5yr | 1-2yr | 8-10yr | 10-15 yr | 5yr | 8=10ye | 25-30yr | 2-3 years |
| 4 | Life span of seed | 4-5yr | 1-2yr | 2-3yrs | 5yr | 4-5 yrs | 1-2 yr | 4-5yr | 2-3years |
| 5 | Protection from rodents | Yes (30-45cm above) | Yes | Yes, hanging from roof | Yes | Yes | Yes | Not | Yes |
| 6 | Temperature control | Yes | Yes | Yes | Not | Not | Yes | Yes | Yes |
| 7 | Humidity control | Not | Yes | Yes | Not | Not | Yes | Yes | Not |
| 8 | Efficiency of seed protectants | Neem, Ash, Cow urine/wood resin | Neem, Cow urine | Neem, Ash | Neem, Ash, cow urine | Neem, Ash, Cow urine/wood resin | Neem, turmeric, lemon, walnut, Cow urine | Neem, cow dung plaster | Neem, cow dung plaster |
| 9 | Handling | Easy | Easy | Easy | Easy | Easy | Easy | Difficult | Difficult |
| 10 | Cost | Low | Low | Medium | Medium | Low | Low | Medium | Medium |

**Table 1b.** Details of the modern seed or grain storage structures used in the study.

|  | **Parameters** | **Pusa bin** | **PAU bin** | **Hapur bin** | **Baked clay bin** | **Metal bins** | **Udaipur bin** |
| --- | --- | --- | --- | --- | --- | --- | --- |
| 1 | Strength of material | Mud, brick, cement | Iron metal | Metal, rubber | Mud, bricks, cement | Steel, Alu. | Coaltar drums |
| 2 | Capacity | 10-30q | 2-15q | 2-10q | 7q | 10q-100q | 1-3q |
| 3 | Life span of structure | 8-10yrs | >15yrs | >15yrs | 5-6yrs | >15-20years | 4-5 years |
| 4 | Life span of seed | 4-5 | 5yrs | 4-5 yrs | 2-3yrs | 5-7 years | 1-2years |
| 5 | Protection from rodents | Yes | Yes | Yes | Yes | Yes | yes |
| 6 | Temperature control | Yes | Yes | Yes | Yes | Yes | yes |
| 7 | Humidity control | Yes, 700-gauge poly. sheet | Yes | Yes | Yes (poly sheet) | Yes | yes |
| 8 | Efficiency of seed protectants | yes | Yes | Yes | Yes | Yes | yes |
| 9 | Handling | Easy | Easy | Easy | Easy | Difficult | Easy |
| 10 | Cost | Medium | High | High | Medium | High | Medium |

**Table 2a.** Assigned scores to traditional seeds/grains storage structures based on selected parameters.

|  | Parameters | Kothi | Gummi | Khalda | Sandook | Kuthar | Matka | Brick structure | Underground structure |
| --- | --- | --- | --- | --- | --- | --- | --- | --- | --- |
| 1 | Strength of material | 3 | 1 | 3 | 3 | 3 | 1 | 5 | 1 |
| 2 | Capacity to store | 3 | 3 | 1 | 5 | 7 | 1 | 7 | 7 |
| 3 | Life span of structure | 3 | 1 | 5 | 3 | 3 | 5 | 7 | 1 |
| 4 | Life span of seed/grain | 3 | 1 | 3 | 3 | 3 | 1 | 3 | 1 |
| 5 | Protection from rodents | 1 | 1 | 1 | 1 | 1 | 1 | 3 | 3 |
| 6 | Temperature control | 3 | 3 | 3 | 1 | 1 | 3 | 3 | 3 |
| 7 | Humidity control | 3 | 3 | 3 | 1 | 1 | 3 | 3 | 3 |
| 8 | Efficiency of plant protectants | 3 | 0 | 1 | 3 | 3 | 3 | 3 | 3 |
| 9 | Handling | 1 | 1 | 1 | 1 | 1 | 1 | 0 | 0 |
| 10 | Cost | -1 | -1 | -3 | -3 | -1 | -1 | -3 | -3 |

**Table 2b.** Assigned scores to modern seeds/grains storage structures based on selected parameters.

|  | Parameters | Pusa bin | PAU bin | Hapur bin | Baked clay bin | Metal bins | Udaipur bin |
| --- | --- | --- | --- | --- | --- | --- | --- |
| 1 | Strength of material | 1 | 7 | 7 | 5 | 7 | 7 |
| 2 | Capacity to store | 7 | 5 | 5 | 5 | 7 | 3 |
| 3 | Life span of structure | 5 | 7 | 7 | 3 | 7 | 3 |
| 4 | Life span of seed/grain | 3 | 3 | 3 | 1 | 5 | 1 |
| 5 | Protection from rodents | 5 | 5 | 5 | 3 | 5 | 3 |
| 6 | Temperature control | 5 | 5 | 5 | 3 | 5 | 3 |
| 7 | Humidity control | 5 | 5 | 5 | 3 | 5 | 3 |
| 8 | Efficiency of plant protectants | 5 | 5 | 5 | 3 | 5 | 5 |
| 9 | Handling | 1 | 1 | 1 | 1 | 0 | 1 |
| 10 | Cost | -5 | -5 | -5 | -3 | -5 | -3 |

- I. Strength of material: Mud and bamboo=1, leather/wood=3, concrete and bricks=5, metal sheet=7 - II. Capacity to store: upto 50kg=1, upto 5q=3, 5-15q=5, >15q=7 - III. Life span of structure: upto 2 years=1, 2-5 years=3, 5-10years=5, >10years=7 - IV. Life span of seed/grain: 2 year=1, upto 5year=3, >5years=5 - V. Protection from rodents: low=1, medium=3, high=5 - VI. Temperature control: low=1, medium=3, high=5 - VII. Humidity control: low=1, medium=3, high=5 - VIII. Protectants: not used=0, used of botanicals or chemicals=1, use of both botanicals and chemicals=3 - IX. Handling: Easy=1, difficult=0 - X. Cost: low=-1, medium=-3, high=-5

### AHP-TOPSIS Technique

The weights for the parameters were calculated by utilizing AHP. The AHP is one of different methods used for the analysis of complicated decisions or multi criteria decision making (Saaty 2008). This method is based on pair-wise comparison and provides scenario checking ability to the researchers and thus helps classifying goals and ways in one complicated environment. The weights assigned to each parameter were used to compute weighed normalized decision matrix. Whereas, TOPSIS method selects the best alternatives which have the nearest distance to the Positive Ideal Solution and farthest distance to the Negative Ideal Solution (Momeni et al. 2011). Further, Euclidean distance was used to compute the distances between each alternative and both the positive-ideal and the negative-ideal solutions. Based on the distance, closeness coefficient is computed to determine the ranking (Taylan et al. 2015). A good account of TOPSIS methodology is given in recent literatures (Yang et al. 2011, Nagpal and Kumar 2012, Iç and Yurdakul 2010).

The steps of hybrid AHP-TOPSIS methodology adopted in this paper are as follows:

Step 1: In this step, decomposition of the decision problem into different elements according to their common characteristics and the formation of hierarchical model were done.

Step 2: Two types of pair wise comparisons are made in the AHP. The first one was between the factor pairs within the same hierarchical level which involves analyst inputs of relative importance ratings based on the pair wise comparative ratings. We refer to Saaty (1996) for the details of pair-wise comparison scale used in the study. The factors weights are computed with following equations (Eq. hereafter) and used in the final hierarchical merit aggregation process.

The matrices of pair-wise comparisons (Eq. 1) are obtained.

In this matrix, the element 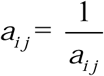 and thus, when i=j *a* _*i j*_ =1. Every element in an upper level is used to compare with respect to the elements in the level below. This work is done by pair-wise comparison two by two and through dedicating numeral scores which shows priority and majority between two decision elements.

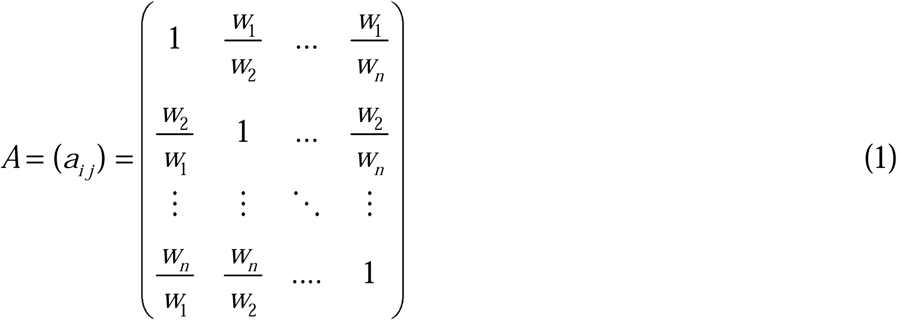

Step 3: The traditional eigen vector method, is the weight vector and is the goal for present study. It helps in measuring the consistency of the referee’s preference arranged in comparison matrix. The Consistency Index (CI) measures the degree of logical consistency among pair-wise comparisons and was used in same manner as defined previously (Saaty 1996) using (Eq. 2):

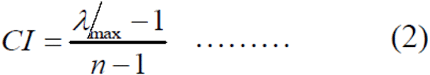

Where, *n* is the number of existing items in the judgment matrix problem.

Consistency Ratio (CR) indicates that the amount of allowed inconsistency i.e. (0.1 or 10%). The score higher than 0.1 indicate that the comparisons are less consistent and smaller than 0.1 means the comparisons are more consistent. CRs above 0.1 means the pair-wise comparison should be revisited or revised again. It is calculated using the following formula of Eq. 3:

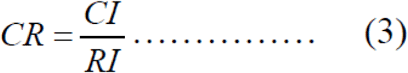

Random Index (RI) for matrices of order (n) 1 to 12 has been as per previous report, for reference see (Saaty 1996).

For AHP, the aggregated final priorities of the alternatives were calculated. Based on that, the weights were constituted and respective weights were used for further analysis. We have selected eight traditional storage structures and six modern storage structures and classified all these on ten parameters.

Step 4. Establish a decision matrix which represents all the alternatives and criteria. The matrix format D can be concisely written as (Eq.4):

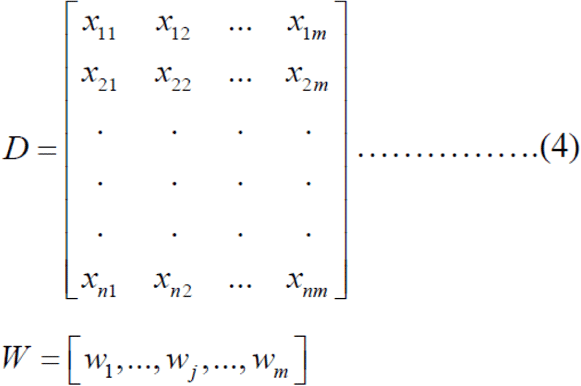

where, *w*_*j*_, (*j*=1,2,…m) is the weight for each criteria (selection parameters for seed storage structures) and *x*_*ij*_(i=1,2,…n ; j=1,2,…m) is the score of i^th^ alternative under j^th^ criteria.

Step 5. The criteria weights are computed by pair wise comparisons matrix employing AHP. Step6. The normalized decision matrix is obtained by employing vector normalization technique. The computation procedure of the element *r*_*ij*_is given as (Eq. 5):

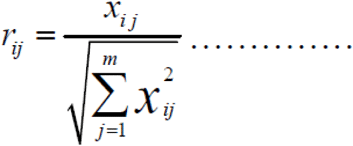

Step 7: The weighted normalized decision matrix can be computed by multiplying each row (*r*_*ij*_) of the normalized decision matrix with its associated attribute weight *w*_*j*_. The weighted normalized value *v*_*ij*_is calculated as below (Eq. 6):

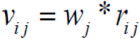

Step 8: The ideal solution (A^+^) is made of all the best performance scores and the negative-ideal solution (A^-^) is made of all the worst performance scores at the measures in the weighted normalized decision matrix. The mathematical expression is written as follows (Eq. 7).

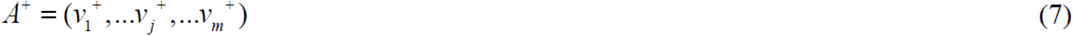

Where,

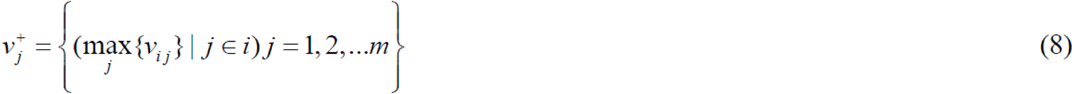

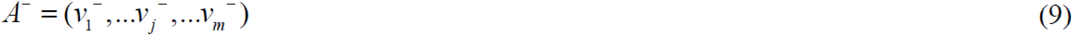

Where

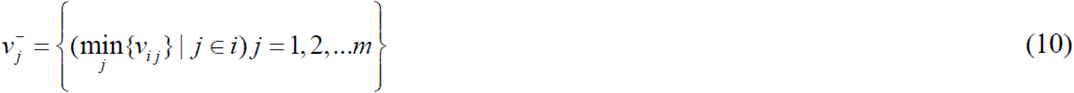

Step 9: The Euclidean distance can measure the distance of an alternative i to the ideal solution. Separation of each alternative from the positive ideal solution (S _i_ ^+^) is then given by Eq. 11.

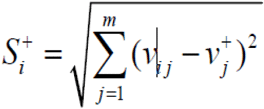

Similarly, separation from the negative ideal solution (S _i_ ^-^) is then given by Eq. 12.

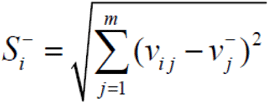

Step 10: The weight (T _i_ ^+^) is computed as (Eq. 13)

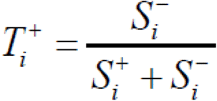

Based on the weight (T _i_ ^+^) final ranking is assign. When weight is close to 1, the alternative is regarded as ideal; and when weight is close to 0, the alternative is regarded as non-ideal. The AHP-TOPSIS hybrid framework for selecting ideal seed storage structures are depicted in Figure 2.

**Figure 2.**
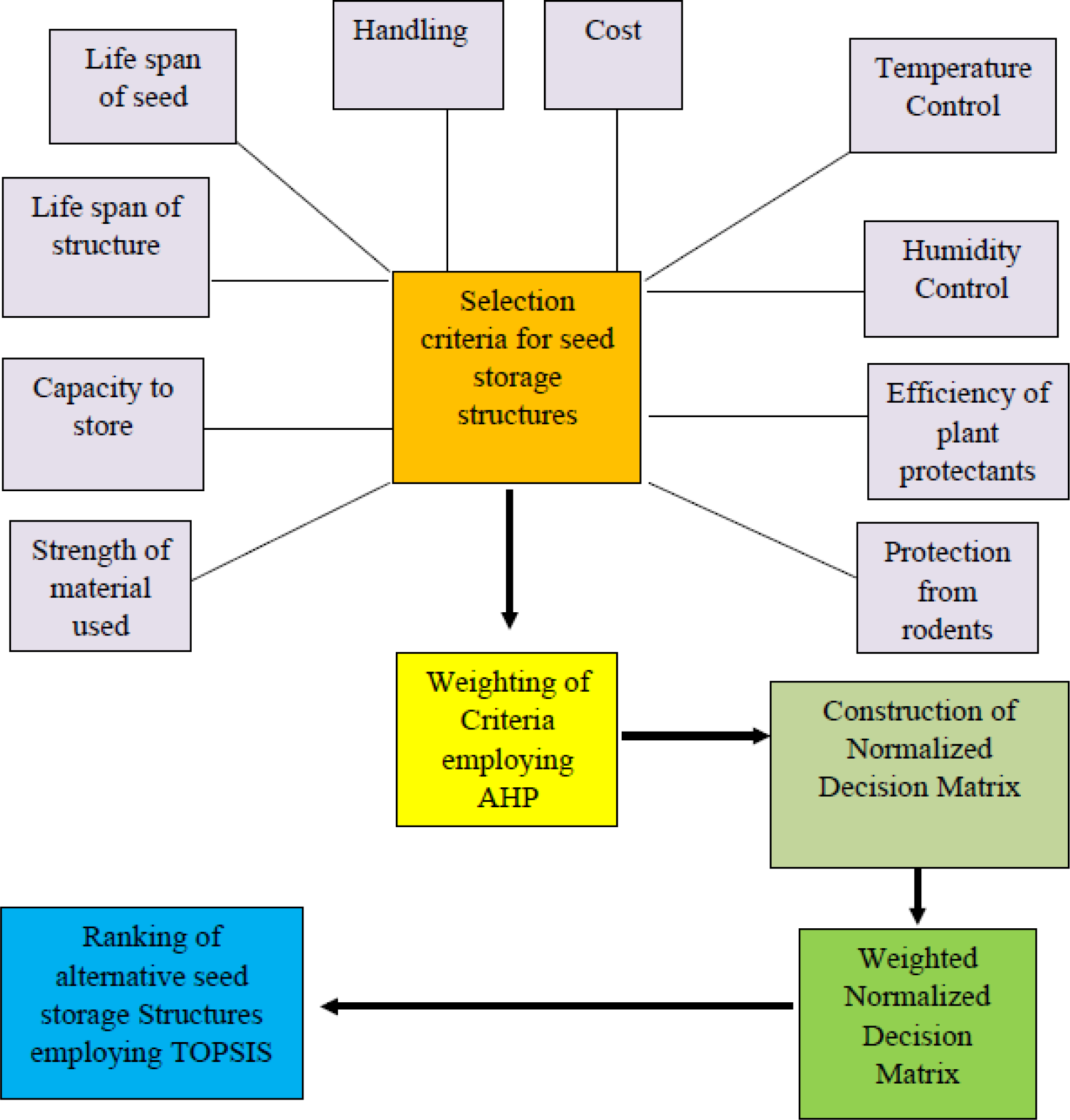
AHP-TOPSIS hybrid framework for selecting ideal seed storage structures.

## Results

### Criteria and Composite Scores of Ideal Seed Storage Structure

The result of survey of specialists based on the AHP methods is depicted in table 3. It indicates the parameters for ideal seed storage structures, their percent weight and eigen vectors. The highest weight was obtained by the parameter temperature control (21 per cent) followed by humidity control (19.6 per cent), protection from rodents (13.5 percent), efficiency of plant protestants used (13 percent) and so on. The consistency index CI was found 0.147 and CR value was calculated as 0.099. These percent score has been further used as weights of respective seed storage parameters. By using the criteria of measurement (Table 3) each storage structures were scored parameter wise (Table 4).

**Table 3:** Selected parameters of seed storage structures on scale of 1 to 7 and their weights.

| Parameter | Measurement (assigned value) | Weights (out of 1) |
| --- | --- | --- |
| Temperature control (C1) | low=1, medium=4, high=7 | 0.2105 |
| Humidity control (C2) | low=1, medium=4, high=7 | 0.196 |
| Efficiency of plant protectants used (C3) | not used=1, used of botanicals or chemicals=4, use of both botanicals and chemicals=7 | 0.130 |
| Protection from rodents (C4) | low=1, medium=4, high=7 | 0.135 |
| Strength of material (C5) | Mud and bamboo=1, leather/wood=3, concrete and bricks=5, metal sheet=7 | 0.115 |
| Capacity to store (C6) | Up to 50kg=1, up to 5q=3, 5-15q=5, >15q=7 | 0.084 |
| Life span of structure (C7) | Up to 2 years=1, 2-5 years=3, 5-10years=5, >10years=7 | 0.046 |
| Life span of seed (C8) | Up to 2 year=1, up to 5year=4, >5years=7 | 0.037 |
| Handling (C9) | Easy=7, difficult=1 | 0.021 |
| Cost (C10) | low=7, medium=4, high=1 | 0.021 |

Note: CI value 0.147; CR value 0.099

**Table 4.**
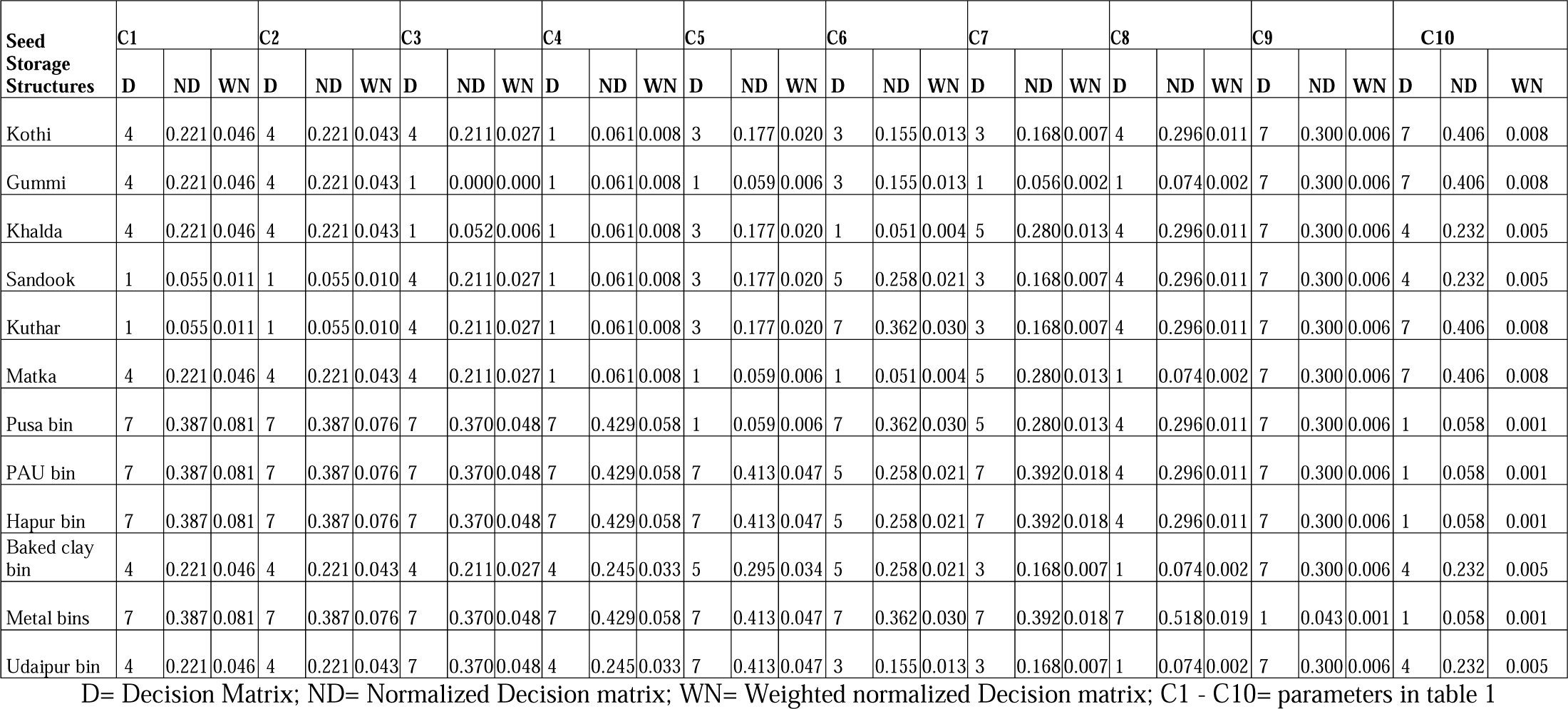
Decision matrix, normalized decision matrix and weighted normalized decision matrix of the seed/grain storage structures.

### Rank of The Storage Structures Based on Composite Score Value

Taking after the AHP TOPSIS methodology, the decision matrix described in Step 4, is made and presented in Table 2. Results indicated that each storage structures scored parameter wise, on a scale from 1 to 7, where 1 is the worst rating and 7 the best, respectively. By applying Eq. 4 as appeared at Step 6 of AHP-TOPSIS methodology, the decision matrix is normalized and the weighted normalized matrix is computed (Table 4) by utilizing Eq. 5. Following the Step 8, the ideal and negative ideal solutions (PIS and NIS) are determined, as shown in Table 5. Then, with Eqs. (10) and (11), the distance from PIS and NIS are calculated respectively, as are shown in table 5. Eventually, the ranking was calculated and the score of each storage structure is calculated and graphically presented in figure 3. According to the analysis, metal bin was estimated as the best seed storage structure with 0.933 score followed by PAU bin and Hapur bin with 0.899 score. The Pusa bin ranked third with 0.740 score followed by Udaipur bin (0.582). All these modern structures scored more because of better provision for temperature control, humidity control, seed protectants and protection from rodents, which have been found to be important parameters for safe storage of seed storage as per the expert judgments and previous studies (Gao and Hailu 2013, Naveenaa et al. 2017, Tefera et al. 2011). In addition, among all modern structures, metal bin was also preferred because of its strength, huge capacity (10-100 quintal) and long-life span (15-20 years). Metal based structures provide comparatively, air-tight module, which do not let pest and pathogens to enter the structure. Moreover, air-tight nature inhibits the growth of any pest and pathogen present from the beginning, due restriction imposed by limited supply of oxygen within the structure (Gao and Hailu 2013, Kumar and Kalita 2017, Tefera et al. 2011, Abass et al. 2014). Among the traditional storage structures brick structure was rated highest (0.549) because of better strength of material, temperature control, and protection from rodents followed by Baked clay bin (0.522), Underground structure (0.469) Kothi (0.424), Matka (0.390), Khalda (0.368), Gummi (0.335), Kuthar (0.272) and Sandook (0.245). From the data it can be inferred that Brick structure and Metal bin are suitable for all the parameters except handling as the handling in these is somewhat less convenient than others. Pusa bin, PAU bin and Hapur bin is suitable for all the parameters except handling and cost. Whereas Gummi, Udaipur bin, Kothi, Khalda and Matka are suitable for handling although the other parameters are not so competitive. Apart from handling, the cost of these modern structures is quite higher than the traditional storage structures. Underground structure, Kuthar, Sandook and Baked clay are found to be suitable for humidity control, protection from rodents and cost. In most of the traditional storage structure, physical parameters fluctuate with surrounding weather conditions, which is responsible for the deterioration of the stored grains (Abass et al. 2014, Chattha et al. 2016). Whereas, the major reasons behind the superiority of concrete based designs over other traditional structures has been due to better provisions for temperature and moisture conditions (Abass et al. 2014).

**Figure 3.**
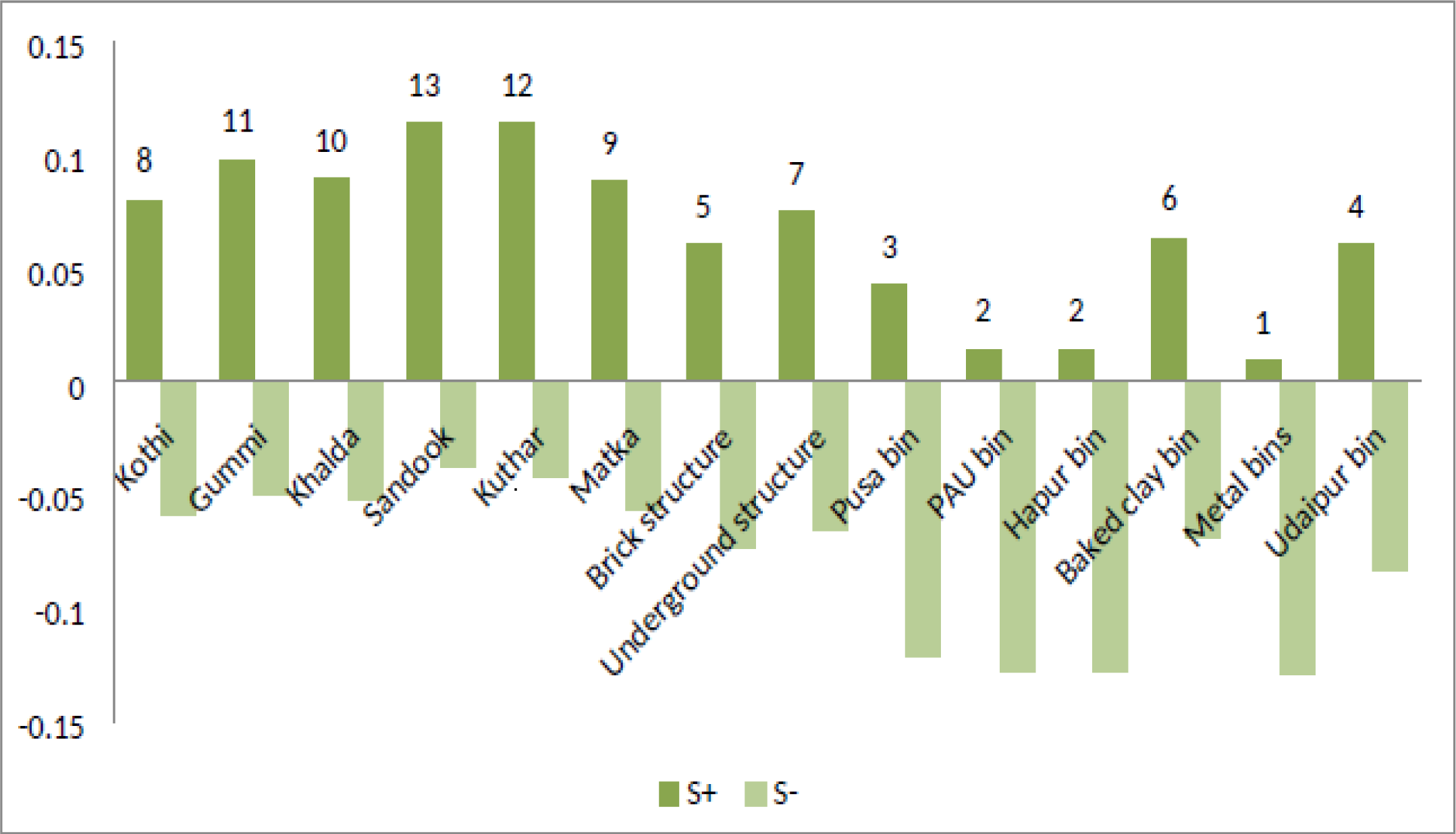
Graphical display of ranks based on distance from positive Ideal Solution and Negative Ideal Solution.

**Table 5:** Positive and Negative ideal Solutions.

|  | C1 | C2 | C3 | C4 | C5 | C6 | C7 | C8 | C9 | C10 |
| --- | --- | --- | --- | --- | --- | --- | --- | --- | --- | --- |
| $A^+$ | 0.081 | 0.076 | 0.048 | 0.058 | 0.047 | 0.030 | 0.018 | 0.019 | 0.006 | 0.081 |
| $A^-$ | 0.011 | 0.011 | 0.000 | 0.008 | 0.006 | 0.004 | 0.002 | 0.002 | 0.001 | 0.011 |

### Proposed Extension Studies for The Improvement of Traditional Structures

From the analysis, it is evident that except brick structure, none of the other traditional structures is appropriate for safe storage of the grains. Having said that, it is also of relevance that, in terms of certain parameters (like C1, C2, C3 and C8) few of them are either equal or have out-competed the modern ones. Hence, in light of these findings, we need to make the statement that, all modern storage structures are not as good as have been documented (Mishra et al. 2012, Nagpal and Kumar 2012). These findings also strengthen the notions, that traditional structures were developed around specific ecological conditions along the line of ingenious crop germplasm as reported in some previous studies (Karthikeyan et al. 2009, Mann et al. 2016) and to a large extent, their diminishing significance could be attributed to changed agriculture methods post green revolution and failure of local tribes to improvise and continuing improvement of their features.

Agricultural technologies are ‘scale neutral’ but not ‘resource neutral’ and from perspective of small farmers, parameters like ease of handling and associated cost cannot be ignored and therefore research and extension studies should give them due priority. Hence, to link marginal farmers, extensions studies directed towards improvement of existing structures in their local settings could also be taken up. Under the provisions of Food and Agriculture Organization (FAO), such extension studies have already been conducted in the neighbouring country Nepal (Ganesh Kumar 1992). As at engineering level, considerable resemblance could be figured out among the grain storage practices between farmers from India and Napal, so taking lead from our analysis and these findings we like to propose following improvements in storage technologies practices by Indian farmers.

- ***Temperature control***: In that study use of poly-lined bags or polythene sacks inside the storage bin have been found to provide an extra layer of insulation and found to secure the structure from the difference between internal and external environments. Likewise, the traditional Indian seed storage containers such as kothi, sandook, kuthar could be sacked with polythene sheets along all sides and painted with bitumen could provide protection from fluctuating external weather.
- ***Humidity control***: A simple strategy was reported to get rid of dampness and thereby control over humidity fluctuation. In the strategy, structures were placed in the house above to ground and away from the sides/walls of the house, about 30-40 cms. In addition to dampness, the strategy also provide protection from rodents. In another, work, painting the external body with white enamel and use of double lid with clay pot like structure was found to control moisture content efficiently. This strategy could be replicated with similar structure in India called ‘Matka’.
- ***Rodent control***: As described above, controlling moisture also provides protection from rodents. From point of engineering, rodent problem could also be reduced, my placing barrier to their entry. For example, with most of the mud, clay on wooden based.
- Traditional structures like Kothi, Gummi, Sandook, Kuthar this could be achieved by placing strips of metals plates all around the base of these structures.
- ***Handling***: Traditional structure Kothi, comes in different variants, static and foldable, similar strategy could also be approached to improve the handling problem of modern metal bins. With joint efforts from FAO and Nepal government a packable kit of metal has been developed and these packable kits have been reported to work quite efficiently (Ganesh Kumar 1992).
- ***Cost efficiency***: As far as cost is considered, one common strategy could be to link the local venders involved in manufacturing these structures with The Mahatma Gandhi National Rural Employment Guarantee Act (MGNREGA) or to provide these structures on subsidize rates to farmers.

## Discussions

In practice, none of the available structures could guarantee complete protection against all pests, how well designed the structure is it is always prone to infestation without use of protectants (Kumar and Kalita 2017, Tefera et al. 2011). Arguably, a storage structure should adhere to these minimum structural necessities so as to provide a certain acceptable level of protection (Tefera et al. 2011). All these traditional storage structures are widely used in villages due to low cost, indigenous knowledge associated with it and easy handling although as per as storage capacity and lifespan of these traditional structures are at the rear end. Most proponents of traditional structures argue about the sustainability and cost benefits of the design for poor community [18, 16]. However, we believe these parameters need further evaluation, as cost benefits and sustainability go hand-in-hand and decide the adoption of a new technology or adherence to the old one. Efficient use of storage technology prevents post-harvest food losses thereby reducing wastage of food and resources used to produce them from very beginning to their final destination in the hands of consumers. All, of these saving provides benefits in form of more sustainable crop production system as well as more socio-economic benefits to poor communities.

India, losses a considerable amount of its produce during storage period which in part could be attributed to the poor seed storage infrastructure and lack of knowledge among farm community (Singh 2010, Mann et al. 2016). Thus, for country like India, it is imperative to understand the usefulness, science, technicality and limitations imposed by the design of storage structures in light of above parameters. Further, we believe that supplying storage facilities to beneficiaries is not the only solution as has been stated (Nagpal and Kumar 2012), rather, we should also try to strengthen the storage facilities available with the farming communities (Malek et al. 2017), as these structures have their own benefits. For instance, underground structures are found to be suitable for humidity control, protection from rodents and cost, thus provisions for better temperature control could enhance their efficiency. Such provisions in traditional structures must be sought, to promote the adaptation of better post-harvest management at the level of the most resource less.

The study proposed a methodology to identify the qualitative and quantitative parameters to be considered for seed storage structures and ranked them based on quantitative scores. Combining and comparing qualitative parameters with quantitative parameters particularly in case of multi criteria decision making is really difficult because of lack of perfect scale of reference. These multi-criteria decision-making methods offers a comparatively transparent way to test the significance of policy related objectives (Vera-Montenegro et al. 2014a and b, Yang et al. 2011). Furthermore, applications of these methods are advantageous in terms of data collection and evaluation, because they are based on qualitative data collected from viewers and/or experts. Industrial application of multi-criteria methodology is widely reported in the literature, including few studies related to agricultural production systems (Vera-Montenegro et al. 2014a and b, Alphonce 1997). However, these studies only included quantitative parameters in the process, neglecting qualitative attributes. In addition, cases related to seed storage structures comparison are not reported. AHP, which was developed and initialised by T.L. Saaty in the 1970’s (Saaty 1987), is among the most widely used techniques in participatory planning for establishment of weights on the criteria and evaluation of plans with respect to each objective (Kangas and Kangas 2005). An alternative to AHP is the Technique for Order Preference by Similarity to Ideal Solution (TOPSIS). TOPSIS, which was developed by Hwang and Yoon in 1981. TOPSIS is based on the concept that the chosen alternative should have the shortest distance to the positive-ideal solution and the longest distance from the negative-ideal solution (Hwang and Yoon 1981). Several authors have reported the use of combined AHP and TOPSIS in multiple criteria problems; where AHP is used to assign weights to the criteria followed by TOPSIS analysis to calculate the final ranking of the alternatives (Gao and Hailu 2013). However, AHP-TOPSIS has not yet been introduced to Post-harvest storage orientated problems.

Our research provides support for the combination of valuation techniques employed to obtain reliable estimates of attribute values for complex goods. Combining AHP and TOPSIS seems to be a promising alternative to AHP alone and to TOSIS. The result showed that the combination of AHP and TOPSIS is practically easy to implement into agriculture extension studies.

## Conclusions

The aim of the study was to evaluate different storage structures based on selected criteria, in a view to support decision making. The study figured out the important parameters for designing effective storage system across different level of storage. The study will help in decision making as well as guide farm community and government agencies in redesigning the available structures based on the parameters highlighted. Further, study would set a paradigm for successful evolution and development of multi-criteria decision making using AHP-TOPSIS hybrid model in post-harvest management studies.

## Acknowledgement

Authors are highly thankful to the end-users of the seed storage structures for providing helpful information while collecting and analysing the data and also to the directors of the respective institutes for providing scientific facility to conduct the research.

